# PLK1-mediated phosphorylation of PHGDH reprograms serine metabolism in advanced prostate cancer

**DOI:** 10.1101/2025.05.21.655274

**Authors:** Xiongjian Rao, Derek B Allison, Robert M Flight, Penghui Lin, Daheng He, Zhiguo Li, Yanquan Zhang, Ruixin Wang, Chaohao Li, Jianlin Wang, Xinyi Wang, Jia Peng, Ka Wing Fong, Qing Shao, Chi Wang, Eunus S. Ali, Hunter NB Moseley, Xiaoqi Liu

## Abstract

Metabolic reprogramming is a hallmark of cancer, enabling tumor cells to meet their increased biosynthetic and energetic demands. While cells possess the capacity for de novo serine biosynthesis, most transformed cancer cells heavily depend on exogenous serine uptake to sustain their growth, yet the regulatory mechanisms driving this metabolic dependency remain poorly understood. Here, we uncover a novel mechanism by which Polo-like kinase 1 (PLK1), often overexpressed in prostate cancer, orchestrates a metabolic shift in serine and lipid metabolism through the phosphorylation of phosphoglycerate dehydrogenase (PHGDH), the rate-limiting enzyme of the serine synthesis pathway (SSP). We demonstrate that PLK1 phosphorylates PHGDH at three specific sites (S512, S513, S517), leading to a marked reduction in its protein level and enzymatic activity. This downregulation of SSP forces cancer cells to increase their reliance on exogenous serine uptake via the ASCT2 transporter, which, in turn, fuels the biosynthesis of lipids, including sphingolipids essential for tumor growth and survival. Targeting the SSP, serine uptake, or downstream lipid biosynthetic pathways may offer promising therapeutic avenues in PLK1-high advanced cancers.

## Introduction

Cancer cells are known for their remarkable ability to reprogram their metabolism to meet the demands of rapid growth and survival under nutrient-poor conditions (Finley, 2023; Hanahan & Weinberg, 2011; Lien & Vander Heiden, 2019; Martínez-Reyes & Chandel, 2021). This metabolic flexibility enables tumor cells to adapt by altering pathways involved in glucose, amino acid, and lipid metabolism (Faubert *et al*, 2020). One of the critical metabolic pathways involved in this process is SSP, which contributes to purine nucleotide biosynthesis, redox balance, and phospholipid production, thus playing an essential role in sustaining tumor growth (Locasale *et al*, 2011; Mattaini *et al*, 2016). PHGDH, the first and rate-limiting enzyme of SSP, has been reported to be overexpressed in multiple cancers, such as breast cancer, melanoma, renal cell carcinoma and non-small cell lung cancer (Possemato *et al*, 2011; Snell, 1984; Teisseire *et al*, 2025; Zhang *et al*, 2017), where it contributes to aggressive phenotypes and resistance to therapy (Mullarky *et al*, 2016; Yoon *et al*, 2023). However, in contrast to its upregulation in certain cancers, PHGDH expression is downregulated in some tumor types, particularly in metastatic cancers due to its metabolic heterogeneity (Rossi *et al*, 2022). Studies have shown that lower PHGDH levels are linked to increased metastatic potential, as the downregulation of de novo serine biosynthesis may enhance the invasiveness of cancer cells by promoting reliance on exogenous serine and other metabolic adaptations (Geeraerts *et al*, 2021; Labuschagne *et al*, 2014). This dual role of PHGDH in different cancers presents a paradox that remains unresolved and highlights the need for a deeper understanding of the regulatory mechanisms that govern serine metabolism along with dependency on serine uptake in cancer.

In parallel, PLK1, a key regulator of mitotic entry and spindle assembly (Conti *et al*, 2024; Gelot *et al*, 2023; Parashara *et al*, 2024), has emerged as a critical player in cancer progression. PLK1 is frequently overexpressed in a variety of malignancies, including prostate cancer, breast cancer, and colorectal cancer (Strebhardt & Ullrich, 2006; Takai *et al*, 2005; Weichert *et al*, 2004), where its high expression correlates with poor prognosis and enhanced tumor growth (Maire *et al*, 2013). While traditionally associated with cell cycle regulation, PLK1 was reported to regulate the pentose phosphate pathway (PPP) (Ma *et al*, 2017). However, despite this insight into its role in PPP, the broader implications of PLK1’s involvement in metabolic reprogramming, beyond cell cycle control, remain largely unexplored. Interestingly, PLK1’s role is context-dependent, as it can inhibit progression in some cancers, adding complexity to its function in tumor biology (de Carcer *et al*, 2018; Raab *et al*, 2018). This dual role of PLK1, coupled with its involvement in metabolic pathways like serine metabolism, highlights a critical gap in understanding how PLK1 orchestrates metabolic shifts in cancer.

While evidence suggests that PLK1 plays a role in cancer metabolism, its specific impact on serine metabolism, particularly in advanced prostate cancer (Gao *et al*, 2019; Reina-Campos *et al*, 2019), remains unclear. To address this gap, we investigate the interplay between PLK1 and serine metabolism, focusing on the regulatory axis between PLK1 signaling and the SSP. Through an integrative approach combining molecular biology, metabolomics, and functional assays, we aim to elucidate how PLK1 reprograms serine metabolism to drive cancer progression.

Our findings not only deepen our understanding of prostate cancer biology but also uncover novel mechanisms underlying cancer’s dependency on serine uptake, highlighting metabolic vulnerabilities that can be exploited for precision therapy. By unraveling this molecular crosstalk, we aim to develop new therapeutic strategies targeting PLK1-mediated metabolic pathways in advanced cancers.

## Results

### PLK1 promotes sphingolipid biosynthesis in prostate cancer

PLK1 is frequently overexpressed in almost all cancers (Fig. EV1A), especially in advanced prostate cancer (Fig. EV1B and Fig. 1A) and showing poor prognosis (Fig. EV1C). The progression of prostate cancer is often associated with aberrant lipid metabolism (Masetti *et al*, 2022; Wang *et al*, 2022), and targeting lipogenesis could be an effective therapeutic strategy for castration-resistant prostate cancer (CRPC) (Zadra *et al*, 2019). To investigate how PLK1 regulates the lipid metabolism to benefit advanced prostate cancer, we examined PLK1 level by western blotting (WB) in normal prostate epithelial cell line RWPE-1 as well as prostate cancer cell lines and then overexpressed PLK1 in LNCaP cell line, which has a relatively lower expression level of PLK1 (Fig. 1B). Next, we performed lipidomic analysis for the extracted lipids from the PLK1-transformed LNCaP cells (LNCaP-PLK1) and control LNCaP cells (LNCaP-CTL) using direct infusion Fourier transform mass spectrometry (DI-FTMS). After removing one outlier, the lipids from the remaining 29 samples (Table EV1) were assigned and classified into major lipid categories: fatty acyls (FA), glycerophospholipids (GP), polyketides (PK), prenol lipids (PR), and sphingolipids (SP). A correlation matrix analysis between these lipids shows high correlation clusters within each lipid category as expected, but also one significant band of high cross-correlation between a large group of glycerophospholipids with a much smaller group of sphingolipids (Fig. 1C). The nominal sums comparison of PLK1/Control is shown in Fig. 1D, and statistical analysis shows that sphingolipid abundances are significantly increased in LNCaP-PLK1 vs LNCaP-CTL at an adjusted p-value of 0.0005, while glycerophospholipids are significantly decreased at an adjusted p-value of 0.05 (Fig. 1E). To determine how PLK1 leads to the increase of sphingolipids, we analyzed the sphingolipid metabolism pathways. As it is known that serine and palmitoyl-CoA are involved in the de novo synthesis of sphingolipids (Fig. 1F), we hypothesized that PLK1 regulates serine metabolism, palmitoyl-CoA metabolism and sphingolipid synthesis. Since there are four major sources of serine (de novo serine synthesis, extracellular serine uptake, serine synthesis from glycine, and autophagy/micropinocytosis) (Newman & Maddocks, 2017; Tsun & Possemato, 2015) and one major source of palmitoyl-CoA (Fig. 1F), we examined the PLK1-dependent regulation of de novo serine synthesis, the serine-glycine transition, de novo lipid (palmitoyl-CoA) synthesis, and sphingolipid metabolism by western blotting (WB). Interestingly, results show that the enzymes of the de novo serine synthesis pathway (SSP) are significantly downregulated upon PLK1 overexpression (OE) (Fig. 1G). These results suggest that PLK1 promotes sphingolipids production in advanced prostate cancer via unexpectedly downregulating SSP.

**Figure 1.**
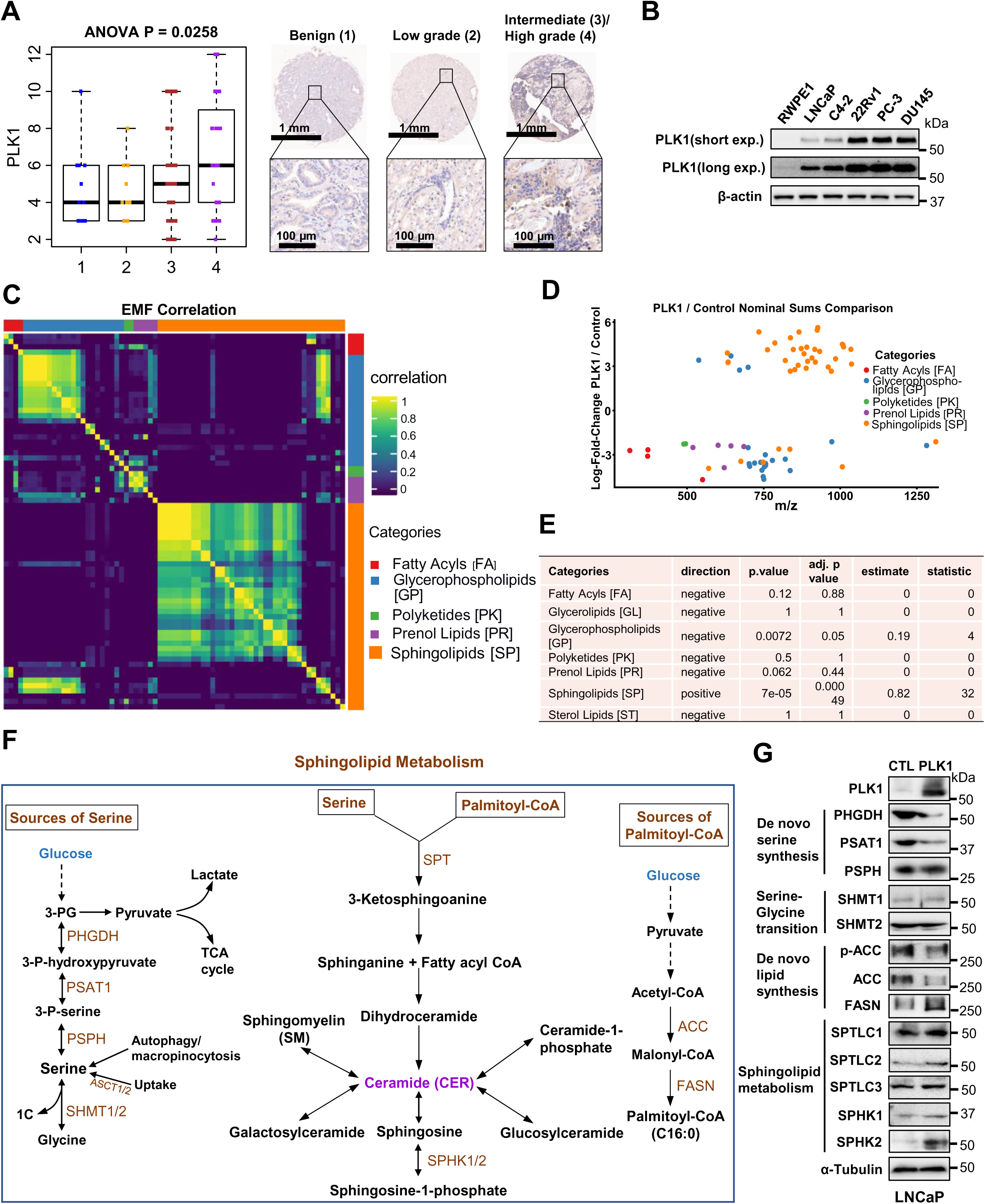
PLK1 promotes sphingolipid metabolism. (A). The ANOVA analysis of PLK1 TMA score and representative images of PLK1 expression in the benign, low grade, intermediate/high grade prostate cancer. Scale bars, 1mm and 100 µm. (B). The PLK1 expression in the prostate cancer cell lines. (C). Elemental molecular formula (EMF) EMF-EMF correlation heatmap. (D). EMF log2 fold-change in PLK1 over Control plotted by M/Z. Points are colored by their voted lipid Category, with “multiple”, “not_lipid” and “not_categorized” removed. (E). Binomial test statistics for each lipid category. (F). Schematic diagram showing the pathways related to sphingolipid metabolism. (G). Immunoblotting of enzymes involved in de novo serine synthesis, serine-glycine transition, de novo lipid synthesis, and sphingolipid metabolism in control and PLK1 overexpressed LNCaP cells.

### PHGDH is downregulated by PLK1

We further confirmed the downregulation of SSP in advanced prostate cancer cell lines (Fig. 2A) and advanced prostate cancer patient-derived xenografts (PDXs) (Fig. 2B), which show the downregulation of PHGDH, the first and rate-limiting enzyme of SSP. We further found that the expression of PHGDH varies across TCGA cancers (Fig. EV2), indicating downregulating SSP is cancer type dependent. To confirm the decrease of PHGDH in advanced prostate cancer, we performed the tissue microarray (TMA) of human prostate samples which showed lower level of PHGDH in advanced prostate cancer (Fig. 2C) and immunohistochemistry (IHC) staining for PLK1 overexpressed mouse prostate cancer samples which showed lower level of PHGDH (Fig. 2D). To further confirm that PLK1 OE leads to the decrease of PHGDH, we co-transfected GFP-tagged PHGDH with different concentrations of HA-tagged PLK1 and found that PHGDH indeed diminishes with the increase of PLK1 (Fig. 2E). To confirm that PLK1 OE leads to the decrease of PHGDH in prostate cancer, we overexpressed PLK1 in LNCaP, C4-2 and 22Rv1 cells, and found that PLK1 OE leads to the decrease of PHGDH (Fig. 2F). To determine if knocking down (KD) PLK1 in advanced prostate cancer cells, such as DU145, leads to the upregulation of PHGDH, we knocked down PLK1 in both C4-2 and DU145 cells, and observed an increase in PHGDH (Fig. 2G). To exclude the mitotic effect of PLK1 and the effects of knockdown PLK1 leading to cell cycle arrest, we set up Tet-inducible PLK1 knockdown cell lines, C4-2-Tet-shPLK1 and DU145-Tet-shPLK1. The PHGDH protein level increased upon doxycycline induction in both C4-2-Tet-shPLK1 cells (Fig. 2H) and DU145-Tet-shPLK1 cells (Fig. 2I). We then KD AR in 22Rv1 cells and overexpressed AR in PC3 cells, which are AR-null and found that PHGDH is decreased upon both AR KD and AR OE (Fig. EV3A,B). We also found PHGDH is decreased in enzalutamide-resistant MR49F cells but increased in enzalutamide resistant C4-2R cells compared to their enzalutamide-sensitive counterparts (Fig. EV3C). Androgen deprivation, an effective therapy for the early-stage prostate cancer, led to an increase in PHGDH in LNCaP cells (Fig. EV3D). Additionally, we also want to know if PLK1 OE in other cancer cell type leads to the decrease of PHGDH, so we overexpressed PLK1 in lung cancer cell line A549 and found the decrease of PHGDH and no change of PSAT1 (Fig. 2J). We also knockdown PLK1 and overexpressed the constitutively active PLK1 T210D (TD) in sarcoma cell line U2OS, and found PHGDH increased upon PLK1 KD, while PHGDH decreased upon TD overexpression (Fig. 2K). These results indicate that PHGDH regulation by AR is not consistent, and we can conclude that PHGDH is regulated by PLK1 rather than AR in prostate cancer.

**Figure 2.**
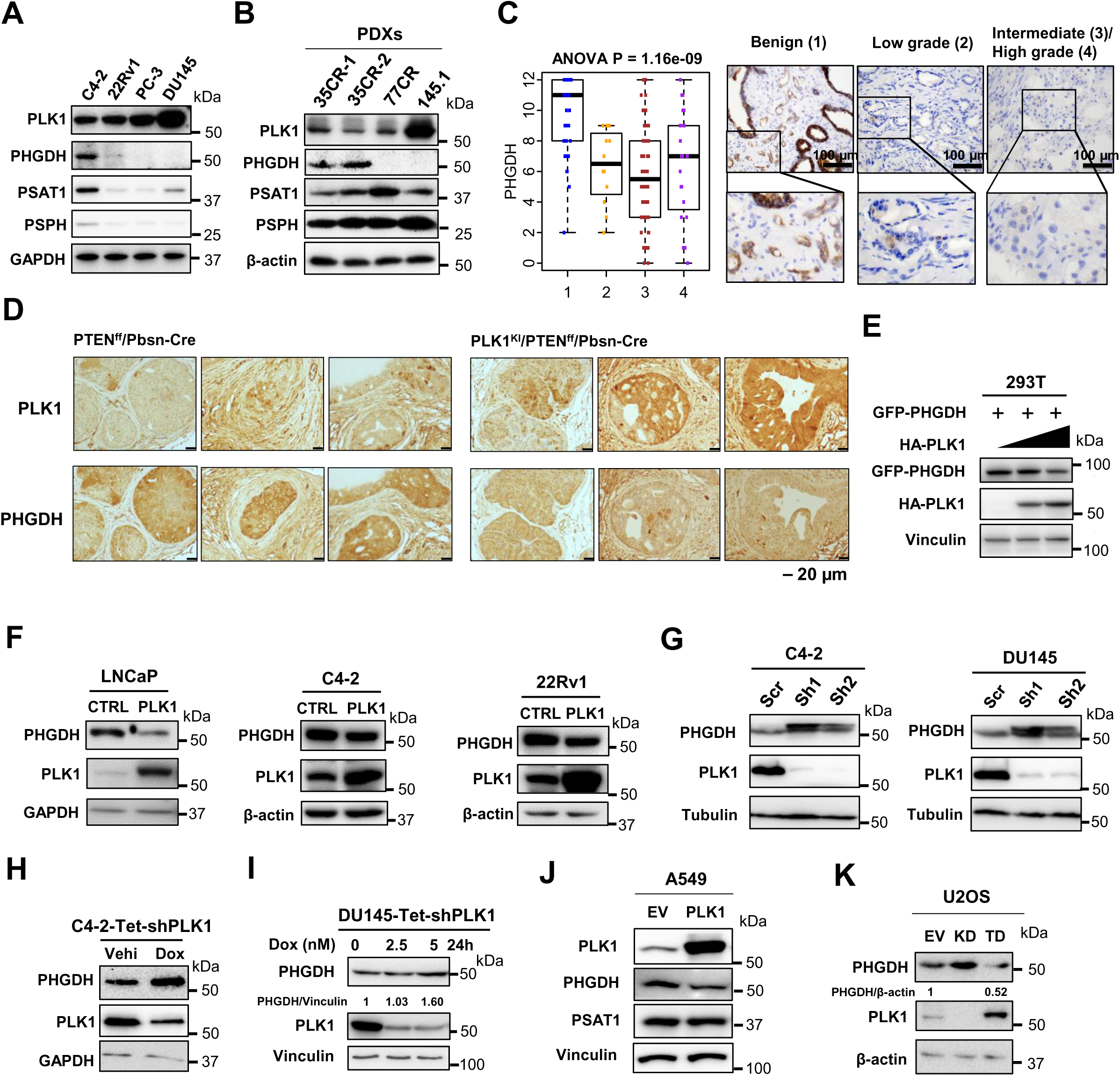
PLK1 downregulates PHGDH in cancer. (A). The expression of enzymes from de novo serine synthesis in prostate cancer cell lines. (B). The expression of enzymes from de novo serine synthesis in prostate cancer PDXs (LuCaP35CR-clone1, LuCaP35CR-clone2, LuCaP77CR, LuCaP145.1). (C). The ANOVA analysis for PLK1 TMA score and representative images of PHGDH expression in the benign, low grade, intermediate/high grade prostate cancer. Scale bars, 100 µm. (D). IHC analysis of PLK1 and PHGDH for prostate tissues from mice with indicated phenotypes. Scale bars, 20 µm. (E). PHGDH level is decreased with the increasing dose of PLK1 expression. GFP-PHGDH and HA-PLK1 co-transfected 293T cells were subjected to IB. (F). PLK1 overexpression (OE) decreases the level of PHGDH. LNCaP, C4-2 and 22Rv1 cells (WT or PLK1-OE) were subjected to immunoblotting (IB). (G). Knockdown (KD) of PLK1 increases the level of PHGDH. C4-2 and Du145 cells (control (scr) and PLK1-KD (sh1/2)) were subjected to IB. (H). Downregulation of PLK1 increases the level of PHGDH. Tet-inducible PLK1 KD C4-2 cells by vehicle (Veh) and 5 nM doxycycline (Dox) were subjected to IB. (I). Downregulation of PLK1 increases the level of PHGDH. Tet-inducible PLK1 KD Du145 cells by doxycycline were subjected to IB. (J). PLK1 overexpression (OE) decreases the level of PHGDH. A549 cells (WT or PLK1-OE) were subjected to immunoblotting (IB). (K). Downregulation of PLK1 increases the level of PHGDH and constitutively active PLK1 (T210D) overexpression (TD) decreases the level of PHGDH. U2OS cells (empty vector (EV), PLK1 knocking down (KD) or PLK1-TD) were subjected to immunoblotting (IB).

### PHGDH is phosphorylated by PLK1

Next, we investigated whether PLK1 binds to and phosphorylates PHGDH via the kinase activity of PLK1. We co-transfected different forms of Flag-tagged PLK1 (WT, K82M, T210D) and PHGDH into 293T cells and immunoprecipitated by flag antibodies to examine the binding interaction between PLK1 and PHGDH. Interestingly, we found that PLK1 bound to PHGDH (Fig. 3A). Moreover, we conducted an in vitro kinase assay and discovered that PHGDH can be phosphorylated by both CDK1 and PLK1 (Fig. 3B). To identify the phosphorylation sites of PHGDH by PLK1, we initially performed an in vitro kinase assay with cold ATP (^31^P-ATP) and analyzed the phosphorylated PHGDH protein using LC-MS/MS. The MS results predicted the phosphorylation sites located between AA 500-520 (Table EV2) with a false positive AA100-110, which we confirmed using the PHGDH fragment (AA1-300) (Fig. 3C). To further map the phosphorylated sites, we conducted a kinase assay by mutating the serine (S) or threonine (T) to Alanine (A) within the AA500-520 (Fig. 3D). We found that the 2A (S512, S513) and 3A (S512, S513, S517) mutations significantly blocked the phosphorylation (Fig. 3E). Thus, we conclude that PHGDH is phosphorylated by PLK1 at multiple sites S512, S513 and S517, which was also confirmed using the online phosphorylation sites prediction website https://gps.biocuckoo.cn/wsresult2.php?p=7, and these sites are consensus among species (Fig. 3F). To investigate the effect of PHGDH phosphorylation by PLK1, we transfected different forms of PHGDH (WT, 3 Aspartic Acid (D) mutation, 3A mutation) into HeLa cells and treated the cells with protein synthesis inhibitor cycloheximide (CHX). Notably, the 3A mutation stabilized PHGDH, whereas the 3D mutation accelerated its degradation (Fig. 3G). Since PHGDH is a metabolic enzyme, we examined whether phosphorylation affects its enzyme activity. Strikingly, we found that PHGDH enzyme activity was significantly reduced upon phosphorylation by PLK1(Fig. 3H), but fairly modest compared with total level of enzyme. Phase transition has been reported as a mechanism of the regulation of enzyme activity regulation (Hardy *et al*, 2024; O’Flynn & Mittag, 2021). We observed that the 3D-mutated PHGDH tends to aggregate into larger dots which may reduce its interaction with substrates compared to the WT form of PHGDH (Fig. 3I).

**Figure 3.**
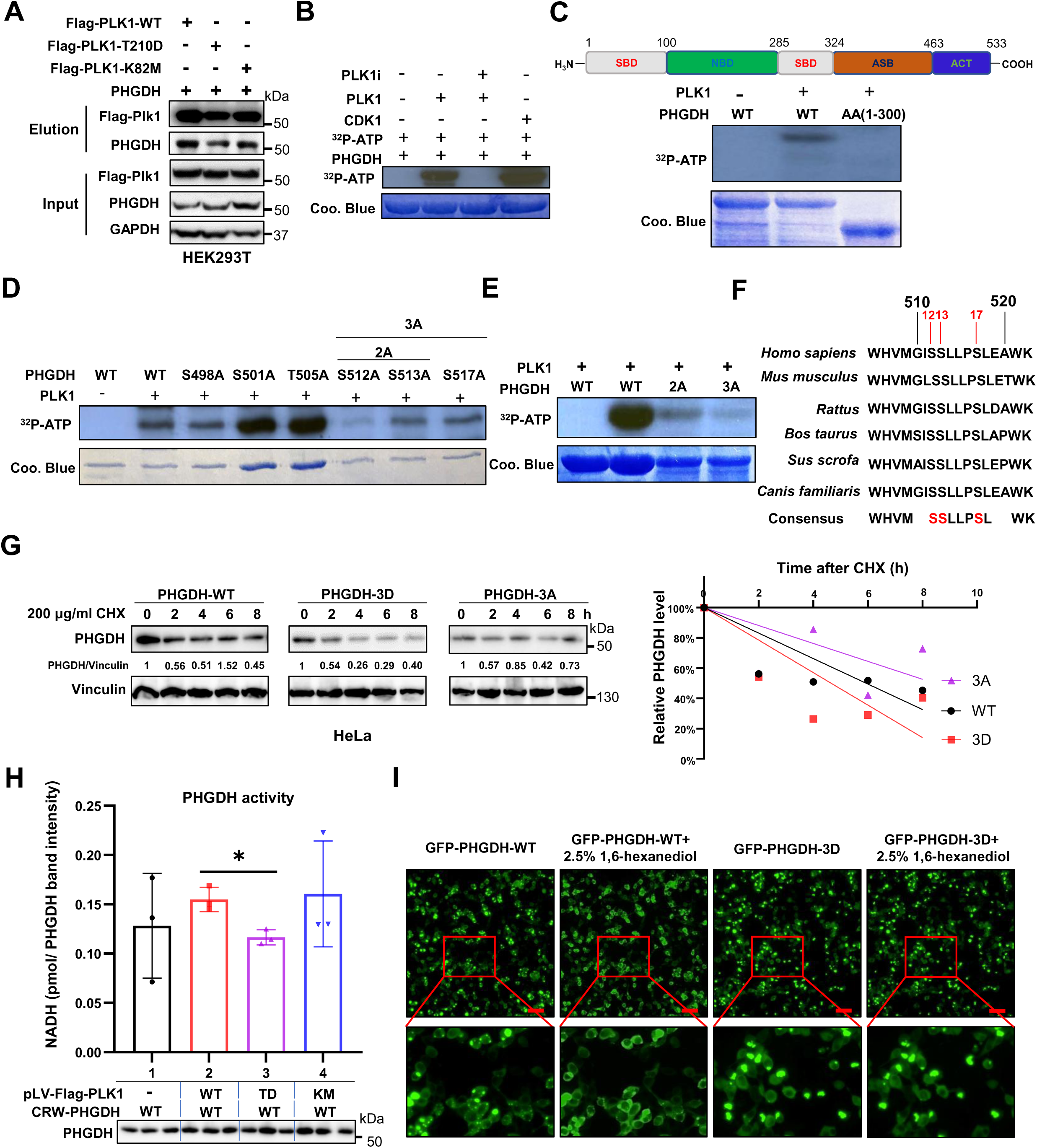
PLK1 phosphorylation of PHGDH leads to its degradation and decrease of enzymatic activity. (A). The binding between PLK1 and PHGDH. Different forms of Flag-tagged PLK1 (WT, T210D, K82M) were co-transfected with PHGDH plasmids in HEK293T cells. The whole cell lysates (WCL) were used for immunoprecipitation (IP) with the M2-Flag beads, followed by western blotting (WB) with targeted antibodies. (B). Kinase assay for purified recombinant WT PHGDH incubated with PLK1, PLK1 as well as PLK1 inhibitor (Onvansertib), and CDK1. (C). Kinase assay for purified recombinant PHGDH (1-300aa) fragments. (D). Kinase assay for purified recombinant PHGDH single A mutants. (E). The kinase assay for PHGDH. Purified His-tagged WT and double/triple A mutants PHGDH were incubated with recombinant PLK1 in the presence of [γ-^32^P]-ATP, followed by SDS-PAGE and film exposure. (F). The consensus amino acid sequencing of PHGDH peptides among species. (G). The protein degradation of PHGDH and mutants. Different forms of PHGDH (WT, 3D, 3A) plasmids were transfected into Hela cells, followed by the treatment of 200 µg/ml CHX. The WCL were examined by PHGDH, and bands were quantified using Image J. (H). The enzyme activity assay for PHGDH. Flag-tagged PLK1 (WT, TD, KM) plasmids were co-transfected with PHGDH (WT, 3A) plasmids. The enzyme activity was measured using a PHGDH enzyme activity assay kit and normalized by the WB band intensity calculation via Image J. Statistical analysis was performed using unpaired two-tailed *t*-tests. Data are shown as means ± s.e.m. *p<0.05. (I). Phase transition of PHGDH protein. GFP-tagged PHGDH (WT, 3D) plasmids were transfected into HEK293T cells. 2.5% 1,6-hexanediol was added into the WT PHGDH and PHGDH-3D plasmids transfected HEK293T cells for 10 min. The cells were imaged under a fluorescence microscope before and after the treatment of 1,6-hexanediol. Bar=50 µm.

To further analyze the effect of phosphorylation on the properties of the PHGDH protein, we conducted large-scale molecular dynamics (MD) simulations and well-tempered meta-dynamics (WT-MeTAD). Fig. EV4A shows the final configurations of the WT and phosphorylated PHGDH in the explicit solvent after a 1000 ns MD simulation at 310 K. The configurations illustrate that the phosphorylation on the three sites induces a change in PHGDH conformation. Fig. EV4B,C show the root mean square fluctuation (RMSF) of Cα atoms in WT and phosphorylated PHGDH during the 1000 ns MD simulations. The RMSF curves indicate that phosphorylation increases the dynamics of PHGDH. We also conducted an analysis on the secondary structure of amino acid residues in certain regions of PHGDH. Phosphorylation dramatically changes the secondary structure of the amino acids in the region 100-200, which is the nucleotide binding domain (NBD), as shown in Fig. EV4D-F. These conformational changes occur in the nucleotide binding domain and thus impact the enzyme activity of PHGDH.

### Prostate cancer expressing high levels of PLK1 tend to import exogenous serine via ASCT2

We found that PLK1 overexpression downregulates SSP via decreasing the protein levels and enzymatic activity of PHGDH. Although serine is classified as a non-essential amino acid, it plays a crucial role as a metabolite for cancer proliferation (Amelio *et al*, 2014; Geeraerts *et al*., 2021). To determine how PLK1 diverts serine metabolism to produce sphingolipids, we setup a Tet-on inducible system to monitor the serine uptake. LNCaP cells were transfected with tet-control (tet-con), tet-PLK1-WT, tet-PLK1-constitively active (tet-PLK1-T210D) plasmids, and stable cell lines were selected using puromycin. These four stable cell lines were cultured in media contained ^14^C-labeled serine to determine serine uptake after doxycycline induction. Our results show that PLK1 overexpression and activation strongly increase serine uptake (Fig. 4A). Since serine is majorly transported into cells through two membrane transporters, ASCT1 and ASCT2 (Conger *et al*, 2024), we performed WB and results show ASCT2 dominates the serine transportation upon PLK1 overexpression (Fig. 4B) and ASCT2 inhibitor V-9302 indeed significantly decreased the serine uptake (Fig. 4A). We also found that ASCT2 is highly expressed in PLK1 highly expressed prostate cancer cell lines, like PC3 and DU145 (Fig. 4C). To confirm the increasing ASCT2 upon PLK1 OE, we overexpressed PLK1 in LNCaP, C4-2 and 22Rv1 cells, and found that PLK1 OE led to the increase of ASCT2 rather than ASCT1 (Fig. 4D), indicating that PLK1 leading to ASCT2 increase is not cell-specific. We also examined the ASCT2 level upon the decrease of PLK1 in the DU145-Tet-shPLK1 cell lines and found ASCT2 decreases with time-dependent treatment of doxycycline (Fig. 4E). To further confirm that highly expressed PLK1 cell lines depends on ASCT2, we treated PC3 and DU145 cells with the PLK1 inhibitor Onvansertib, which causes the decrease of ASCT2 (Fig. 4F). Additionally, our findings show that ASCT2 expression is upregulated in response to PLK1 overexpression, even under conditions of serine deprivation, while the mitochondrial serine transporter SFXN1(Kory *et al*, 2018) was not affected (Fig. 4G). To further test if the decrease of de novo serine synthesis leads to upregulation of ASCT2, we showed that knockdown of PHGDH in C4-2 cells indeed led to the increase of ASCT2, while restoration of PHGDH led to a decrease in ASCT2 (Fig. 4H). We also confirmed that PLK1 OE in A549 leads to the increase of ASCT2 (Fig. EV5A) and KD PLK1 in U2OS led to the decrease of ASCT2 with the TD increasing ASCT2 (Fig. EV5B). The regulation of ASCT2 and sphingolipid metabolism by PLK1 is not associated with the mRNA level of serine transporters and enzymes involved in the sphingolipid metabolism (Fig. EV6A,B). Thus, we conclude that PLK1 decreases de novo serine synthesis by downregulating PHGDH and compensates for the reduced serine pool by upregulating the plasma membrane serine transporter ASCT2, thereby enhancing the exogenous serine uptake.

**Figure 4.**
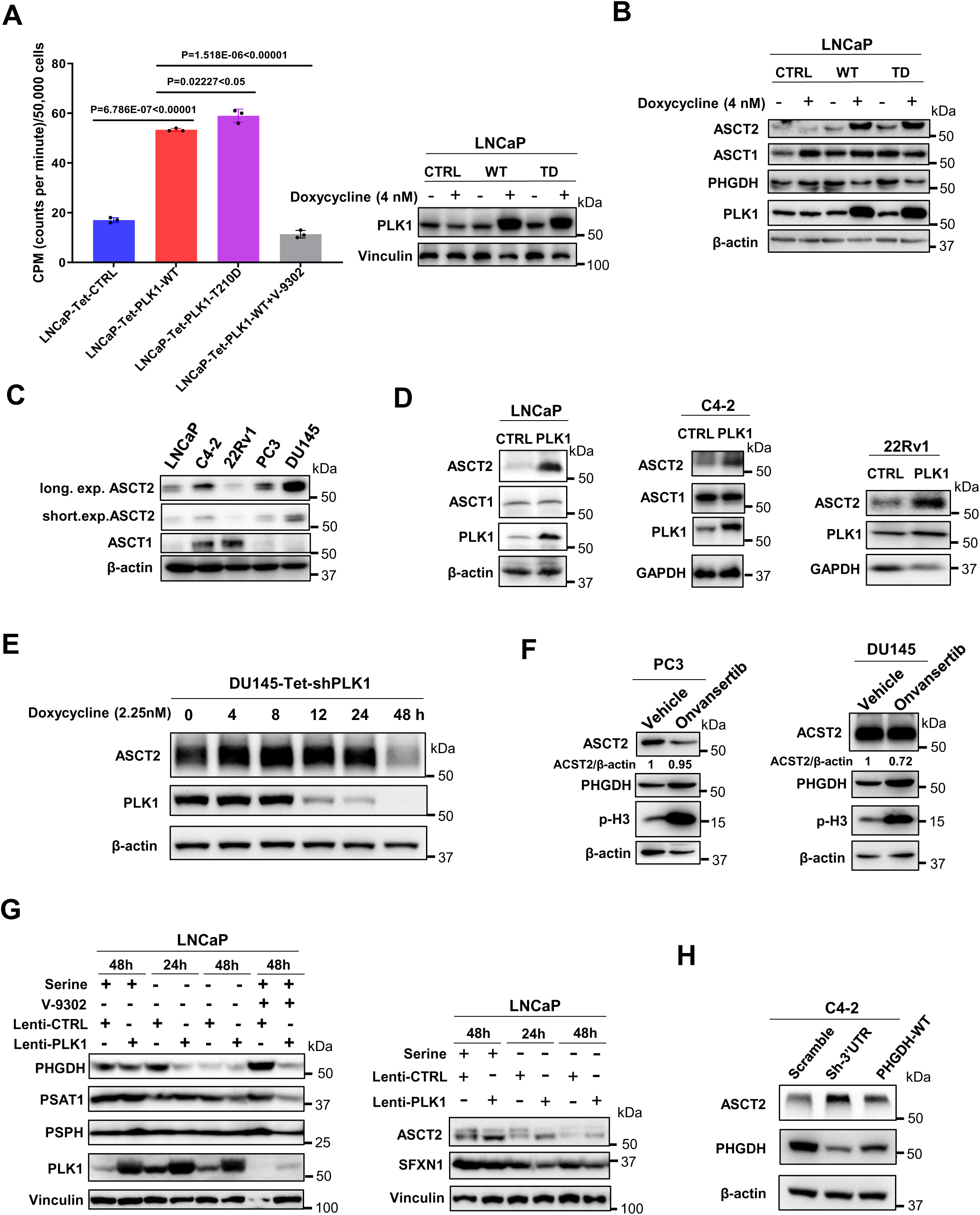
PLK1 promotes exogenous serine uptake via ASCT2. (A). PLK1 overexpression increases serine uptake. The same number (5×10^5^) of LNCaP-Tet-CTL, LNCaP-Tet-PLK1-WT, LNCaP-Tet-PLK1-T210D, and LNCaP-Tet-PLK1-WT cells treated by 5 µM ASCT2 inhibitor V-9302 were seeded onto 6-well plates. Cells were washed and incubated with [^14^C]-serine upon PLK1 expression was induced by 4 nM doxycycline for 24h. Statistical analysis was performed using unpaired two-tailed *t*-tests. (B). ACST1 and ASCT2 expression after PLK1 overexpression in LNCaP cells. (C). ACST1 and ASCT2 expression among prostate cancer cell lines. (D). ASCT2 expression after PLK1 overexpression in LNCaP, C4-2 and 22Rv1 cells. (E). ASCT2 expression after doxycycline induced PLK1 knocking down in DU145-Tet-shPLK1 cells. (F). ASCT2 expression in PC3 and DU145 cells after the treatment of PLK1 inhibitor Onvansertib. (G). The expression of ASCT2 and enzymes involved in SSP upon the absence of serine for 24h and 48h in LNCaP-control and LNCaP-PLK1 cells. (H). ASCT2 expression in the PHGDH knockdown and PHGDH restored C4-2 cells.

### PHGDH phosphorylation by PLK1 causes metabolic reprogramming

To determine the metabolic effects of PHGDH phosphorylation by PLK1, we first setup several cell lines (like C4-2-shRNA Scramble, shRNA PHGDH 3’UTR, shRNA PHGDH 3’UTR-PHGDH WT, shRNA PHGDH 3’UTR-PHGDH 3D, shRNA PHGDH 3’UTR-PHGDH 3A) (Fig.EV7) and then cultured the C4-2-PHGDH WT (WT), C4-2-PHGDH 3D (3D), LNCaP-CTL, and LNCaP-PLK1 cells in the media containing uniformly ^13^C-labeled glucose ([U-^13^C]-Glucose) and deuterium-labeled serine ([2,3,3-^2^H]-serine) to examine glycolysis, TCA cycles, the pentose phosphate pathway (PPP) and nucleotide synthesis. Strikingly, we found that both the PHGDH 3D mutation and PLK1 overexpression affected glycolysis by reducing PEP (Fig. 5A), which is regulated by the enzyme pyruvate kinase M2 form (PKM2). PKM2 was previously reported to be allosterically regulated by serine (Chaneton *et al*, 2012), whose uptake is enhanced by PLK1 overexpression. Our finding that PLK1 leads to the decrease of PHGDH, thus decreasing of SSP in prostate cancer, which is reminiscent of checking inosine monophosphate (IMP), the downstream purine from glucose and one carbon metabolism. We found that the M+1 IMP showed consistent decrease between 3D/WT and PLK1/CTL (Fig. 5B). Additionally, M+1 Glycerol 3-phosphate derived from glucose showed a significant decrease in 3D/WT and PLK1/CTL (Fig. 5C). The PHGDH 3D mutation did not affect the Krebs cycle (TCA cycle) (Fig. EV8A), PPP (Fig. EV8B), synthesis of other nucleotides from glucose (Fig. EV9A), or other glucose-related metabolic pathways (Fig. EV9B,C). We then investigated whether PHGDH phosphorylation by PLK1 affects exogenous serine incorporation into nucleotide synthesis (Fig. EV10A). We confirmed that ^2^H incorporation from ^2^H3-Ser into metabolites involved in the nucleotide metabolic pathways and found that ^2^H incorporation into nucleotides was inconsistent between 3D/WT and PLK1/CTL (Fig. EV10B-D). These results indicate that PHGDH phosphorylation by PLK1 decreases SSP as well as glucose-derived serine into one carbon metabolism, but does not affect the exogenous serine incorporation into one carbon metabolism and the downstream nucleotides.

**Figure 5.**
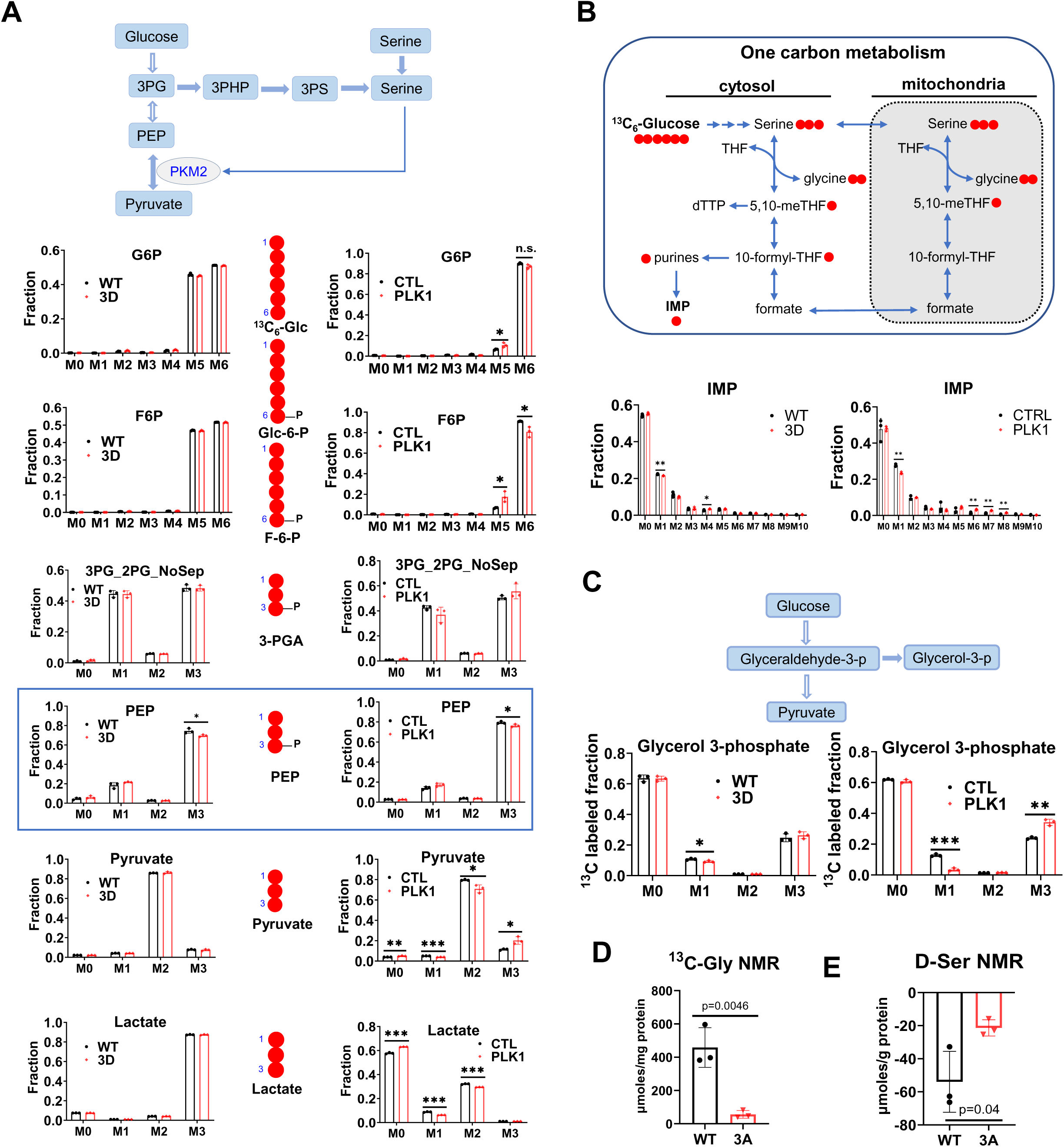
PHGDH phosphorylation by PLK1 causes metabolomic reprogramming. (A). The comparison of labeling glycolytic intermediates using [U]-^13^C-glucose between LNCaP-PLK1 vs LNCaP-Control cells (right) and C4-2-PHGDH-3D vs C4-2-PHGDH-WT cells (left). (B). The comparison of ^13^C incorporation into Inosine monophosphate (IMP) from [U]-^13^C-glucose between LNCaP-PLK1 vs LNCaP-Control cells and C4-2-PHGDH-3D vs C4-2-PHGDH-WT cells. (C). The comparison of glycerol 3-phosphate labelling from [U]-^13^C-glucose between LNCaP-PLK1 vs LNCaP-Control cells and C4-2-PHGDH-3D vs C4-2-PHGDH-WT cells. (D). The 1D ^1^H(^13^C)-HSQC NMR analysis for the glucose-derived glycine production for C4-2-3A and C4-2-WT cells. (E). The deuterium (^2^H) NMR analysis for D-serine (D-Ser) from the media of C4-2-3A and C4-2-WT cells. Statistical analysis was performed using unpaired two-tailed *t*-tests (**A**-**E**). Data are shown as means ± s.e.m. * p<0.05, ** p<0.01, *** p<0.001.

To establish that blocking PHGDH phosphorylation by PLK1 leads to specific metabolic changes, we used NMR to analyze the media and polar fractions from WT, 3D and C4-2-PHGDH-3A (3A) cells. We found that PLK1 overexpression enhanced glucose consumption and led to significant pyruvate production, but not lactate production (Fig. EV11A). Blocking PHGDH phosphorylation with the 3A mutation resulted in no apparent reduction in glucose consumption and significant lactate production compared with WT (Fig. EV11B), while both the reduction in glucose consumption and lactate production were significant compared to 3D (Fig. EV11C). We also examined the excretion of glutamine, alanine and glutamate, which were as expected lower in 3A (Fig. EV11D). The production of glucose, lactate, alanine, glycine, asparagine, succinate, fumarate, and glutamate, among others, decreased after blocking PHGDH phosphorylation by PLK1 using the 3A mutation (Fig. EV11E). The heatmap of RNA-seq and Kyoto Encyclopedia of Genes and Genomes (KEGG) pathways showed the decrease of sphingolipid metabolism and glycine, serine and threonine metabolism pathways after blocking phosphorylation of PHGDH by 3A mutation (Fig. EV12). The consumption of deuterated serine was significantly reduced after blocking the phosphorylation of PHGDH by 3A mutation (Fig. 5D,E). These results not only confirmed our conclusion that PHGDH is phosphorylated by PLK1 leading to decreased activity, but also indicate that PHGDH phosphorylation by PLK1 affects exogenous serine uptake, thus affecting the downstream pathways, especially sphingolipid metabolism, glycine, serine and threonine metabolism.

### PHGDH phosphorylation by PLK1 leads to lipid remodeling

To investigate whether exogenous serine is incorporated into serine-containing lipids, we analyzed deuterium(^2^H) incorporation into lipids from WT and 3A cells. Our analysis revealed that over 37% of the detected lipids among the grand total lipids were labeled with ^2^H derived from serine (Table EV3), indicating that exogenous serine is a critical precursor for lipid biosynthesis in prostate cancer cells. Among these, the most abundant ^2^H-labeled lipid species were triglyceride (TAG, 11.86%), phosphatidylethanolamine plasmalogen (PE-pmg, 9.82%), phosphatidylcholine (PC, 6.99%), phosphatidylethanolamine (PE, 7.89%), phosphatidylcholine plasmalogen (PC-pmg, 2.27%), and phosphatidylglycerol (PG, 1.12%). As these lipids do not contain serine, the deuterium must have arisen from serine metabolism releasing ^2^H, which is then incorporated into the pools of lipid synthesis (Fig. 6A). Higher ^2^H incorporation into serine-containing lipids may reflect incorporation of the exogenous deuterated serine in addition to deuteration of, presumably, the fatty acyl chains. No significant differences in these proportions were observed between WT and 3A cells. Interestingly, among the serine-containing lipids, a much higher proportion of ^2^H labeled species were sphingomyelin (SM), but not phosphatidylserine (PS). More than 98% of the detected sphingolipids were SM in both WT and 3A cells, with the remaining proportion consisting primarily of ceramide (CER) (Table EV4). Blocking PHGDH phosphorylation did not affect ^2^H incorporation into SM, but significantly reduced incorporation into CER, along with some other lipids which are related to ceramide metabolism, like diacylglycerol (DAG), phosphatidylglycerol (PG) and 15:0-18:1(d7) PC 33:1 (PC-18:1) (Fig. 6B).

**Figure 6.**
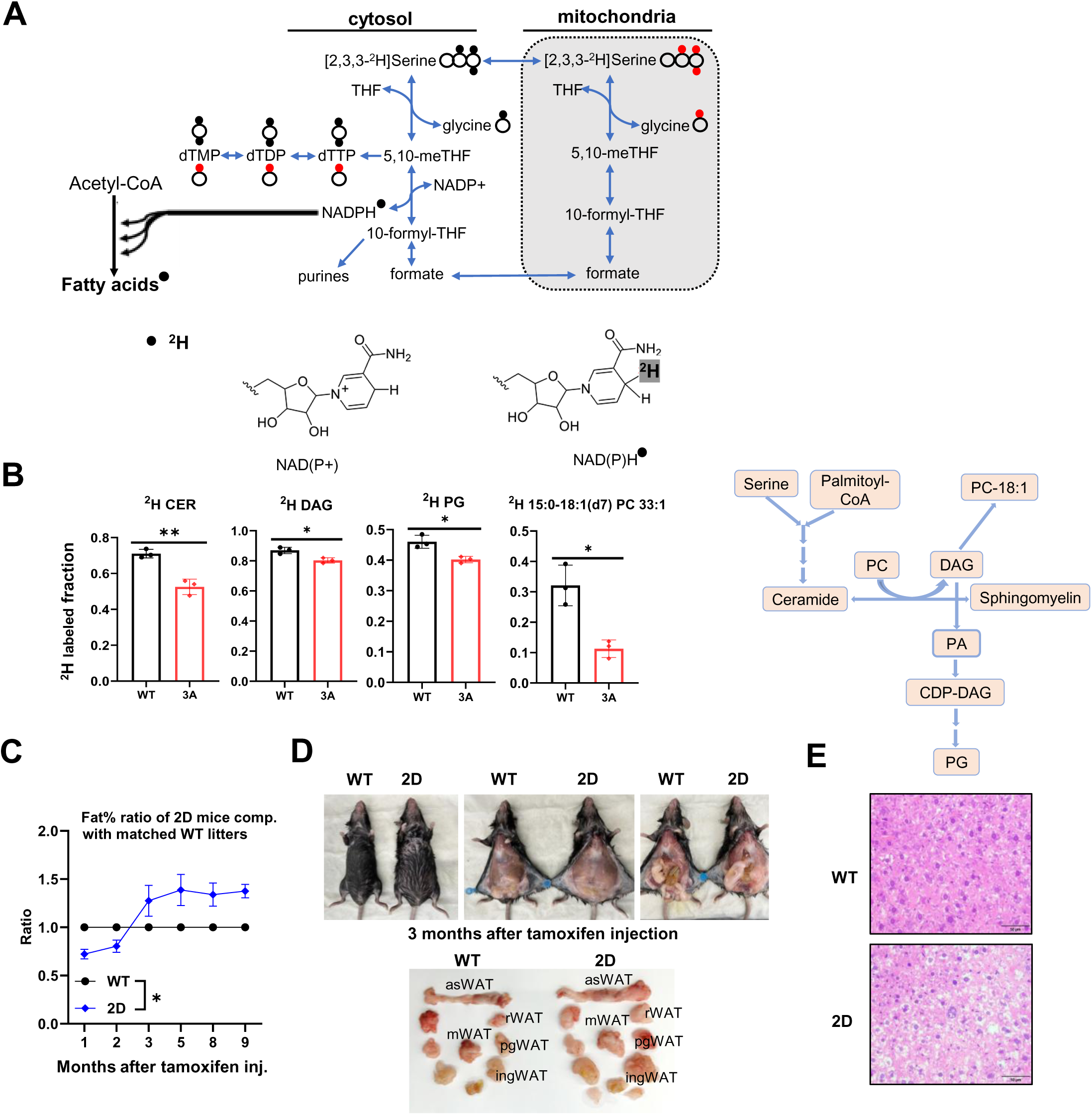
PLK1 phosphorylation of PHGDH affects serine incorporation into lipids and lipid synthesis in vivo. (A). The flow chart of 2H from [2,3,3-2H]-serine incorporation into fatty acids. (B). The [2,3,3-^2^H]-serine incorporation into lipid species in C4-2-PHGDH-WT cells and C4-2-PHGDH-3A cells. Fraction= ^2^H labeled specie/total specie. (C). The fat% ratio (fat/weight of 2D mice**/** fat/weight of WT litter mice) over the time after tamoxifen injection (n>5). (D). Representative images of the dissection of fat tissues from paired WT and 2D mice which are from the same litter. (E). Representative images of H&E staining for the liver from paired WT and 2D mice which are from the same litter.

To assess the physiological consequences of PLK1-mediated phosphorylation of PHGDH on serine metabolism in vivo, we conducted a series of experiments using a mouse model. PHGDH 2D Mice were generated by Taconic, and the mutations were induced by the injection of tamoxifen after crossing the mice with ERT2-Cre mice. The tamoxifen-injected mice were monitored over time using EchoMRI. We found that the 2D heterogenous mice significantly gained fat in both brown and white adipose tissue compared with WT litters (Fig. 6C). We found that the major fat tissues gained by the 2D mice compared with WT mice are anterior-subcutaneous WAT (asWAT), retroperitoneal WAT (rWAT), mesenteric WAT (mWAT), posterior-subcutaneous WAT (psWAT), and inguinal WAT (ingWAT) (Fig. 6D). In comparison to the wild-type mouse, the mutant mouse shows histologic evidence of hepatocyte unrest, which can be seen in the setting of inflammation, toxins, or metabolic disturbances. However, in these histologic sections, there is no increase in inflammation and no evidence of toxic injury to the liver. As a result, these findings could be compatible with a metabolic dysregulation process (Fig. 6E). These in vivo findings further confirmed our study that PHGDH phosphorylation by PLK1 leads to lipid remodeling.

### PLK1-mediated PHGDH phosphorylation and lipid remodeling offer potential therapeutic targets

Metabolic profiling of PLK1-transformed prostate cancer cells revealed significant alterations in serine metabolism in advanced prostate cancers. Specifically, mass spectrometry analysis demonstrated a marked reduction in intratumoral levels of serine and its downstream metabolites, indicating decreased de novo serine synthesis and increased serine uptake. Conversely, there was a pronounced increase in the levels of sphingolipids, derived in part from serine, in PLK1-overexpressing tumors compared to controls.

Collectively, these findings provide compelling evidence that PLK1-mediated phosphorylation of PHGDH regulates serine metabolism, leading to metabolic reprogramming characterized by decreased de novo serine biosynthesis and enhanced sphingolipid (and apparently general lipogenesis) production. These metabolic alterations highlight the therapeutic potential of targeting PLK1-mediated metabolic vulnerabilities in cancer therapies. The combination treatment of PHGDH inhibitor NCT-503 and PLK1 inhibitor Onvansertib effectively block the prostate cancer growth (Fig. 7A). PLK1 overexpressed C4-2 and 22Rv1 cells are more sensitive to ASCT2 inhibitor V-9302 (Fig. 7B,C). PHGDH-3D C4-2 cells are more sensitive to serine or serine/glycine deprivation compared to PHGDH-3A C4-2 cells (Fig. 7D). PLK1 overexpressed 22Rv1 cells are more sensitive to the sphingosine kinase 2 (SPHK2) inhibitor ABC294640 (Fig. 7E). Interestingly, PLK1 highly expressed N2P1 xenografts are more sensitive to PHGDH inhibitor NCT-503 compared to 22Rv1 xenografts (Fig. 7F,G), indicating that tumors usually characterize nutrient-deprived conditions which need serine from both exogenous uptake and the SSP. Thus, inhibiting de novo serine synthesis, hindering sphingolipid metabolism and depriving of serine and glycine may provide therapeutic avenues for advanced prostate cancers (Fig. 7H,I). Collectively, this study demonstrates that the oncogenic PLK1 inhibits PHGDH, and subsequently compensatory serine uptake and changes in sphingolipid metabolism ultimately drive the cellular proliferation and prostate tumor growth. These results further indicate that ASCT2, through exogenous serine intake, is a crucial downstream mediator of PLK1-dependent growth of cells and tumor where PHGDH protein expression is persistently low.

**Figure 7.**
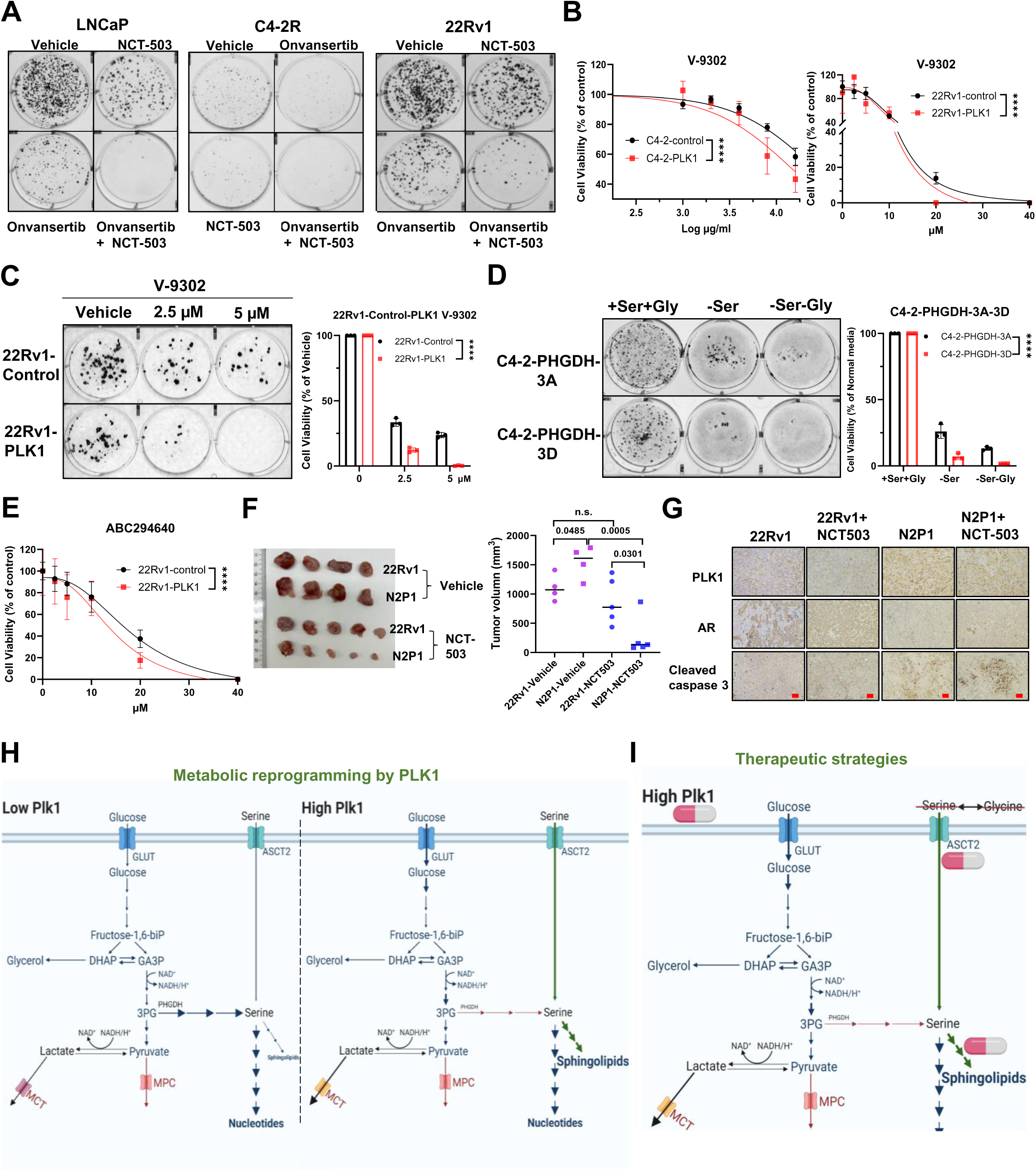
Therapeutic strategies for prostate cancer. (A). Colony formation for LNCaP, C4-2R and 22Rv1 cells after treatment by PLK1 inhibitor Onvansertib, PHGDH inhibitor NCT-503 and the combination of BI6727 and NCT-503. (B). Cell viability assay for C4-2-control/C4-2-PLK1 and 22Rv1-control/22Rv1-PLK1 with the treatment of different concentrations of ASCT2 inhibitor V-9302 for 24h. Statistical analysis was performed using two-way ANOVA. (C). Colony formation and quantification for the 22Rv1-control and 22Rv1-PLK1 cells after treatment by ASCT2 inhibitor V-9302. Statistical analysis was performed using two-way ANOVA. (D). Colony formation and quantification for the C4-2-3A and C4-2-3D cells after the deprivation of serine, glycine, serine and glycine. Statistical analysis was performed using two-way ANOVA. (E). The cell viability of 22Rv1-control and 22Rv1-PLK1 after treatment by SPHK2 inhibitor ABC294640. Statistical analysis was performed using two-way ANOVA. (F). Xenograft images and quantification of 22Rv1 and N2P1 with or without treatment of NCT-503. Statistical analysis was performed using unpaired two-tailed t-tests. (G). IHC analysis of PLK1, AR and cleaved caspase-3 for the xenograft of 22Rv1 and N2P1 with or without treatment of NCT-503. Scale bars, 100 µm. (H). Schematic diagram of metabolic reprogramming by PLK1 in prostate cancer. (I). Schematic diagram of therapeutic strategies Data are shown as means ± s.e.m. * p<0.05, ** p<0.01, n.s., not significant.

## Discussion

Our study reveals a novel regulatory axis in advanced prostate cancer, where PLK1 signaling reprograms serine metabolism, driving tumor growth and progression. PLK1 modulates the activity of PHGDH, the first enzyme in de novo serine biosynthesis, by inducing its phosphorylation and degradation. This destabilization of PHGDH diminishes endogenous serine production, shifting cancer cells toward increased exogenous serine uptake to fuel critical pathways like sphingolipid biosynthesis.

This reprogramming highlights PLK1’s dual role in cell cycle regulation and metabolic control, positioning it as a central orchestrator of cancer metabolism. The downregulation of PHGDH (Fig. 1G and Fig. 2F), a key enzyme often upregulated in other cancer types (Liu *et al*, 2020; Pacold *et al*, 2016; Zhang *et al*, 2023) and non-catalytically expressed lower in cancer dissemination and metastasis (Rossi *et al*., 2022), underscores PLK1’s unique ability to redirect metabolic flux in prostate cancer. Moreover, PLK1-overexpressing cells exhibit an enhanced reliance on extracellular nutrients to sustain their growth, suggesting that both de novo serine biosynthesis and serine uptake are potential therapeutic targets for cancers.

Notably, while PSAT1 and PSPH exhibited no change in expression by PLK1, PHGDH showed a significant decrease in our experimental models of advanced metastatic prostate cancers. This finding provides a distinct example of metabolic diversity in prostate cancers, and the metabolic subtype, as defined by low level of PHGDH and its reliance on serine obtained from the outside of cells.

Metabolomic analysis revealed distinct labeling patterns, including a significant decrease in M+1 Glycerol 3-Phosphate (G3P) (Fig. 5B), which likely originates from the pentose phosphate pathway (Mason *et al*, 2021; Talwar *et al*, 2023), and the dominance of M+2 pyruvate from uniformly labeled ^13^C-glucose (Fig. 5A). These patterns reflect PLK1’s capacity to channel glucose-derived carbons into anabolic pathways rather than merely sustaining energy production.

While the Warburg effect is a hallmark of various cancers, our study shows that PLK1-overexpressing LNCaP prostate cancer cells do not significantly elevate lactate secretion despite enhanced glycolysis (Fig. EV 11A), whereas C4-2-PHGDH-3D cells secret abundant lactate (Fig. EV11C). This cell-specific deviation from the classical Warburg phenotype suggests alternative metabolic fates for pyruvate, likely towards biosynthetic processes rather than energy production via lactate fermentation.

Our unbiased lipidomic screening found out the enhanced sphingolipid synthesis by PLK1 in prostate cancer cell line LNCaP, and we confirmed the significant incorporation of exogenous serine into ceramide (Fig. 6B), which accounts for over 1% of detected sphingolipids (Table EV4) in C4-2 PHGDH mutant cells. While ceramide levels are low, its metabolic messengers (Summers *et al*, 2019) and roles in increasing tumor risk (Li *et al*, 2023) suggest that even minor perturbations in its synthesis could have outsized impacts. Furthermore, phosphorylation of PHGDH by PLK1’s highly expression in tumor which includes both the tumor cells and tumor microenvironment cells amplified the effects on lipid synthesis, which might cause the enhanced fat production in the whole body of mice (Fig. 6C-E).

In conclusion, our findings reveal how PLK1-driven metabolic reprogramming enhances the adaptability of prostate cancer cells, particularly by modulating serine metabolism, glycolysis, and sphingolipid synthesis. These insights open new therapeutic avenues by targeting PLK1-regulated metabolic pathways in advanced prostate cancer. Further investigation into PLK1’s impact on cancer metabolism could lead to the development of novel precision medicine strategies to combat this aggressive disease.

Despite the novel insights presented in this study, several limitations need to be acknowledged. First, the high salt content in samples affected the sensitivity of serine and glycine measurements by Electrospray Ionization Mass Spectrometry (ESI-MS), limiting precise quantification. Second, although we generated phospho-specific antibodies targeting PHGDH at S512 and S513, their insufficient sensitivity, combined with the unclear molecular mechanism behind PHGDH downregulation in response to elevated PLK1 expression, hindered robustly detection of phosphorylated PHGDH. Finally, further studies are needed to assess the effects of a serine/glycine-deficient diet in PHGDH-2D mice and to verify the fat tissue composition via mass spectrometry. These gaps present avenues for future research.

## Materials and Methods

### Cell culture experiments

RWPE-1 cells were cultured in Keratinocyte Serum Free Medium (K-SFM) containing 0.05 mg/ml bovine pituitary extract (BPE) and 5 ng/ml human recombinant epidermal growth factor (EGF). LNCaP, 22Rv1, C4-2B cells were cultured in Roswell Park Memorial Institute 1640 media (RPMI 1640, GIBCO). PC3, DU145, Hela, A549, U2OS and HEK293T cells were cultured in Dulbecco’s Modified Eagles Medium (DMEM). MR49F and C4-2 ER were enzalutamide-resistant cells derived from LNCaP and C4-2 cells, respectively. N2P1 cells were derived from the LuCaP 145.1 patient derived xenograft (PDX). Androgen deprivation therapy (ADT) experiments for LNCaP cells were performed using media supplemented with androgen-free serum, such as charcoal stripped fetal bovine serum (CS-FBS), and dihydrotestosterone (DHT) as control. LNCaP-(CTL, PLK1) cells were selected by puromycin after infection of LNCaP cells with control and PLK1-overexpression lentiviruses. DU145-(CTL, shPLK1) were selected by puromycin after infection of DU145 cells with control and PLK1-knockdown lentiviruses. 22Rv1-(CTL, KD) cells were selected by puromycin after infection of control and AR knockdown lentiviruses in 22Rv1 cells. PC3- (CTL, AR) cells were selected by G418 after infection of control and AR overexpressed lentiviruses in PC3 cells. LNCaP-Tet-(control, PLK1-WT, PLK1-T210D, PLK1-K82M), C4-2-Tet-shPLK1, DU145-Tet-shPLK1 cells were selected by puromycin after infection by the indicated Tet-inducible viruses. Media were supplemented with 10% fetal bovine serum (FBS), 2 mM L-glutamine, 100 U/mL penicillin, and 100 mg/mL streptomycin. The cells were cultured under conditions of 95% air, 5% CO2, 85-95% humidity, 7.2-7.5 pH, 37 °C. Lentiviruses were prepared and used in accordance with the Lentivirus Production Protocols from Addgene.

### Cell lysis, immunoprecipitation (IP) and western blotting (WB)

Cells were lysed using RIPA buffer (20 mM Tris-HCl, 37 mM NaCl, 2 mM EDTA, 1% Triton-X, 10% glycerol, 0.1% SDS, and 0.5% sodium deoxycholate) containing the phosphatase inhibitor (Active Motif, Cat#37492) and protease inhibitor (Sigma, Cat#11836170001) for protein analysis. For immunoprecipitations, cell lysates were incubated with the indicated magnetic beads (anti-Flag M2 magnetic beads, MilliporeSigma) at 4°C for overnight (12h) and washed three times with cold lysis buffer or PBS buffer. Cell extracts and immunoprecipitated proteins were denatured by boiling, separated by SDS-PAGE, transferred to nitrocellulose (NC) or PVDF membranes (Thermo Fisher Scientific), and immunoblotted with the specific antibodies. The membranes were stripped using Restore^TM^ PLUS Western Blot Stripping Buffer (ThermoFisher, Cat#46430) according to the manufacturer’s instructions.

Primary antibodies used include PLK1(MilliporeSigma, Cat#05-844), SHMT1(cSHMT, Santa Cruz,Cat#sc-365203), SHMT2(mSHMT, Santa Cruz,Cat#sc-390641), PHGDH (CST, Cat#66350S and Santa Cruz,Cat#sc-1003177), PSAT1 (Proteintech,Cat#10501-1-AP), PSPH(Peoteintech,Cat#14513-1-AP), ASCT1 (CST,Cat#8442S and Proteintech,Cat#13067-2-AP), ASCT2 (CST,Cat#8057S and Proteintech,Cat#20350-1-AP), ACC(CST,Cat#3676S), p-ACC(CST,Cat#11818S), FASN (CST, Cat#3189S), SPHK1 (Proteintech,Cat#10670-1-AP), SPHK2 (Proteintech, Cat#17096-1-AP), SPTLC1 (Proteintech, Cat#15376-1-AP), SPTLC2 (Proteintech,Cat#51012-2-AP), SPTLC3 (ThermoFisher, Cat#PA5-98858), SFXN1(Proteintech, Cat#12296-1-AP), Flag (MilliporeSigma, Cat#B3111), AR (CST, Cat#19672), PSA (CST, Cat#5365),HA (CST, Cat#3724), GAPDH (CST, Cat#2118L), α-Tubulin (CST, Cat#3873S), β-actin (CST, Cat#4970 and Santa Cruz, Cat#sc-47778), Vinculin (MilliporeSigma, Cat#V4505).

### Immunohistochemistry (IHC) analysis

IHC was performed following the manufacturer’s instructions (vector laboratories). Briefly, tissues were fixed, sectioned, and mounted onto glass slides. Slides were steamed for antigen retrieval, followed by quenching / blocking, incubation with primary antibodies / lectins, secondary antibodies incubation, and application of a tertiary reagent for substrate / chromogen development. After counterstaining, the slides were mounted with coverslips. The images were acquired using a Nikon ECLIPSE Ti2 microscope, capturing at least eight fields of view for each sample.

### Transfection

Plasmids were transfected into 293T cells following the manufacturer’s transfection protocol (Polyplus, jetPRIME). Transfected cells were harvested and lysed in RIPA buffer after 36-48 hours post-transfection for subsequent IP and WB analysis.

### PHGDH enzyme assay

The PHGDH enzyme assay was conducted according to the manufacturer’s instructions (BioVision, Cat#K569-100; Abcam, Cat#ab273328).

### Purification of recombinant proteins

GST-tagged or His-tagged wild type (WT) and mutant PHGDH vectors were cloned and transformed into BL21 E. coli. Protein expression was induced with Isopropyl β-d-1-thiogalactopyranoside (IPTG) for 4 hours at 37°C or approximately 20 hours at 18 °C. Bacterial pellets were centrifuged, resuspended in cold PBS buffer containing lysozyme and NP-40, and then lysed by sonication. The supernatant was incubated with Glutathione Sepharose 4B (GE Healthcare) or cOmplete His-Tag Purification Resin (Roche) at 4°C for overnight (12-16h). Purified PHGDH protein were identified by Coomassie blue staining.

### Kinase assay

The reaction system consisted of 5 µL 10X Kinase buffer (CST, Cat#9802), 1 µL 200ng recombinant PLK1, 1 µL 5µCi ^32^P-ATP (VWR), and purified proteins. The mixture was adjusted to a final volume of 50µL with ddH2O and incubated at 30 °C for 30 minutes. The reacted mixture was then boiled for SDS-PAGE and gels were stained with Coomassie blue. Destained gels were dried using a Hoefer™ GD 2000 Gel Dryer System (Fisher scientific) for 2 hours, and dried gels were exposed to X-ray films for sensitivity analysis.

### Colony Formation Assay

To assess the long-term proliferative potential of prostate cancer cells under various experimental conditions, a colony formation assay was performed. Cells (e.g., LNCaP, C4-2R, or 22Rv1) were seeded at a low density of 500-1000 cells per well in 6-well plates. After allowing the cells to attach overnight, the appropriate treatments (e.g., drug treatments) were applied the following day. The cells were maintained under standard culture conditions (37°C, 5% CO₂) with media changes every 3 days for approximately 10-14 days, until visible colonies formed.

Once colonies became prominent, the media was removed, and the cells were gently washed with phosphate-buffered saline (PBS). The colonies were fixed with 4% paraformaldehyde for 15 minutes at room temperature and stained with 0.5% crystal violet for 30 minutes. The excess stain was removed by washing with distilled water. Plates were air-dried, and colony numbers were counted manually using a stereomicroscope. A colony was defined as a cluster of at least 50 cells. The colony formation efficiency was calculated as the ratio of the number of colonies formed to the number of cells initially seeded.

### Mouse strains

PHGDH-S512D/S513D (PHGDH-2D) mice were generated by Taconic Biosciences. This strain models a phosphorylation-mimetic variant of PHGDH, a key enzyme in serine biosynthesis, enabling investigation of its metabolic or oncogenic roles. ERT2-cre (strain#008463), Pbsn-cre (strain#026662), Pten^flox^ (strain#006440) mice were acquired from The Jackson Laboratory. ERT2-cre allows tamoxifen-inducible Cre recombination for temporal control of genetic manipulation, while Pbsn-cre drives prostate-specific expression if prostate biology is under study. The Pten^flox^ strain enables conditional knockout of this tumor suppressor, a common approach in cancer models. Mice were crossed to generate PHGDH-2D/ERT2-cre^+^ littermates, which were observed or used without major sex-specific differences. Groups in individual experiments were derived from one litter or were sex- and age-matched. Mice were treated constitutively with tamoxifen (75 mg/kg, oral gavage) for 5 days to induce the Cre expression according to the Jackson Laboratory’s instructions. Genotyping was conducted using the ear or tail pieces, following the Jackson Laboratory’s instructions. All mouse experiments were approved by the official Kentucky ethics committee for animal experiments and University of Kentucky (UKy) Animal Care and Use Committee (Protocol no: 2020–3680).

### EchoMRI

The mice which were treated with tamoxifen (allowing time for Cre-induction) were weighed and performed EchoMRI assay for body composition in the UK-COCVD COBRE Mouse Metabolism Core, under procedures approved in IACUC protocol 2016-2337. EchoMRI provides a non-invasive, precise measurement of fat/lean mass, avoiding terminal procedures at this stage. The use of a dedicated core facility ensures standardized, high-quality metabolic phenotyping.

### Tissue microarray (TMA)

Tissue microarrays (TMA) were prepared from a cohort of human prostate biopsies. All tissue fixation, H&E staining, and IHC analysis were performed at the UKy Biospecimen Procurement & Translational Pathology Shared Resource Facility (BPTP SRF).

### Xenograft study

Cells (22Rv1 and N2P1) were subcutaneously inoculated into pre-castrated nude mice (from The Jackson Laboratory) and drug treatment commenced when tumor reached 100 mm^3^ in size. Tumor volume (V = L x W x H/2) were measured every 2 days, and tumors were harvested before reaching 2000 mm^3^. Harvested tumors were weighed, fixed with 4% formaldehyde, sectioned, and subjected to IHC analysis with indicated antibodies.

### Cell viability assay

Cells were seeded at 6000-8000 cells/100 µl/well in 96-well culture plates and allowed to grow overnight. Drug treatment was performed based on compound toxicity, with incubation at 37 °C for 24-72 hours. Cell viability was observed daily under a microscope, and when ∼90% cell death was observed at the higher drug concentration(s), the AquaBluer™ assay was conducted. 100 µl of diluted AquaBluer™ reagent (Boca Scientific) was added to each well using a multi-channel pipettor, and plates were incubated for 4 hours. Fluorescence intensity (RFU) was measured at 540ex/590em using the GloMax Discover Microplate Reader (Promega).

### Bioinformatic analysis

TCGA-PRAD (Prostate Adenocarcinoma) RNA-Seq gene expression data were downloaded from the Genomic Data Commons (https://gdc.cancer.gov) and normalized by TPM (Transcripts Per Kilobase Million). The processed TCGA-PAAD clinical data was downloaded from cBioPortal. Kaplan-Meier curves and logrank tests were used to compare the survival probabilities between PLK1 and PHGDH high and low subgroups by median expression respectively. The Cox proportional hazards model was used to estimate the hazard ratios between the PLK1 and PHGDH high and low subgroups. Additionally, non-parametric Spearman correlation was applied to evaluate the correlation between PLK1 and PHGDH mRNA expressions within tumor samples.

For the identification of phosphorylation sites, we used Biocuckoo computational platform (Wang *et al*, 2020) to predict PLK1-mediated phosphorylation sites of PHGDH.

### L-[^14^C]-serine uptake

Cells (5×10^5^) were seeded into 35mm diameter dishes (or 6-well plates) for 24 hours in triplicate per cell line before L-[U-^14^C]-Serine uptake experiments. LNCaP-control and LNCaP-PLK1 cells were first analyzed to determine if PLK1 overexpression could increase serine uptake. Second, C4-2-control and C4-2-PLK1 cells, 22Rv1-control and 22Rv1-PLK1 cells, DU145-control and DU145-shPLK1 cells were analyzed to further confirm the hypothesis. Third, the following prostate cancer cell lines, LNCaP, C4-2, 22Rv1, PC3, DU145 were used to compare the serine uptake. After 24 h seeding, culture media were removed and cells were carefully washed with pre-warmed Hanks’ Balanced Salt Solution (HBSS) (137 mM NaCl, 1.3 mM CaCl2, 0.5 mM MgCl2, 0.4 mM MgSO4, 5.3 mM KCl, 0.44 mM KH2PO4, 4.2 mM NaHCO3, 0.33 mM Na2HPO4, 5.6 mM glucose, 25 mM HEPES), pre-incubated in 1.0 mL of pre-warmed HBSS at 37 °C for 5 min before adding L-[U-^14^C]-Serine for the uptake experiment. Cells were then incubated at 37 °C for 5 min in 1 mL of HBSS containing 10 μM L-[U-^14^C]-amino acid (1.11 kBq/mL) (PerkinElmer Life Sciences/ VWR). Subsequently, cells were washed carefully three times with ice-cold Na^+^-free HBSS containing 1.0 mM non-radiolabeled amino acid. Then, the cells were lysed with 500 μL of 0.1 N NaOH and mixed with 3 mL of Ultima Gold (PerkinElmer). Radioactivity were measured using a liquid β-scintillation counter (Beckman LS 6000SE). All dishes/plates were placed into protective chambers both in the incubators and hoods. All washed HBSS /cells and used dished/plates/tips/tubes were discarded into radiation-labeled containers. After experiments, all manipulated area (incubator, hood, counter) were sampled using cotton swabs for radioactivity test to make sure the related area is clean.

### Molecular dynamics simulations

Large-scale molecular dynamics (MD) simulations and well-tempered meta-dynamics (WT-MeTAD) were conducted to analyze the effect of phosphorylation on the properties of the PHGDH protein. The initial configuration of a full-length WT PHGDH was predicted using alphaFold2 (Jumper *et al*, 2021) based on the amino acid sequence. The initial configuration of the phosphorylated PHGDH was derived from the WT one using an online post-translational modification server (Margreitter *et al*, 2013). MD simulations were conducted using Gromacs (Abraham *et al*, 2015) with GPU acceleration on the computing facilities at the University of Kentucky.

### Proteomics analysis of phosphorylation

For proteomics analysis of PHGHD phosphorylation by PLK1, an in vitro kinase assay with cold ATP (^31^P-ATP) was performed, followed by enrichment of phosphorylated PHGDH using the High-Selec^TM^ TiO2 Phosphopeptide Enrichment Kit (ThermoScintific, Cat#A32993). The enriched phosphorylated PHGDH was resolved using SDS-PAGE and the gel bands were excised for trypsin digestion. The digested phosphopeptides were fragmented using the Thermo Exploris 240 Orbitrap Mass Spectrometer to identify peptide sequences and potential phosphorylation sites.

### Stable Isotope-Resolved Metabolomics (SIRM) experiments

For SIRM experiments, cells were cultured in 10 cm plates until approximately 70% confluence overnight. The media were then replaced with isotope tracer media for 24 hours. Media samples were harvested at different time points (e.g. 0, 3, 6, 12 and 24h) following the replacement of isotope tracer media containing 10% dialyzed serum, and extracted using cold acetone (final concentration 80% acetone). Cells were quenched with a cold mixture of CH3CN and nanopore water at the ratio of 2:1.5 (v/v). Polar, non-polar, and protein samples were harvested by centrifugation after the addition of chloroform. Residual materials were washed by a cold mixture of chloroform, methanol, butylated hydroxytoluene (BHT) at a of 2:1:1mM ratio and centrifuged to collect remaining polar, non-polar, and protein residues. All samples were lyophilized using a liquid nitrogen (LN2) trap and stored at -80 °C for further nuclear magnetic resonance (NMR) and mass spectrometry (MS) analysis.

### NMR spectroscopy

NMR spectra were recorded at 15°C on a 16.45 T Bruker Avance III spectrometer equipped with a 1.7 mm triple resonance (HCN) cryoprobe.

For ^1^H spectra, lyophilized samples were redissolved in 35 µL deuterated phosphate buffer containing 17.5 nmole d6-2,2-dimethyl-2-silapentane-5-sulfonate (DSS-d6) as an internal reference. 1D ^1^H spectra were acquired using the zgpr pulse sequence with a spectral width of 12 ppm, and acquisition time of 2 s and a relaxation delay of 4 s and 512 transients. The residual HOD signal was suppressed using presaturation with a weak transmitter pulse during the relaxation delay. Under these conditions, the magnetization recovers > 95 % for most signals. Free induction decays were zero-filled to 128 K data points and apodised with a 1 Hz exponential function, Fourier transformed, phased, baseline corrected and referenced to the internal DSS peak. 1D ^1^H(^13^C)-HSQC (heteronuclear single quantum coherence) experiments were recorded with a spectral width of 12 ppm, 1024 transients, a relaxation delay of 1.75 s, and an acquisition time of 0.25 s, during which adiabatic decoupling was applied. ^1^H(^13^C)-HSQC spectra were processed with zero filling to 16,384 data points and apodized with a cosine squared function and a 4 Hz exponential line broadening, Fourier transformed, phased, baseline corrected and referenced to the internal DSS peak.

For deuterium (^2^H) spectra, lyophilized samples were redissolved in 35 µl ^1^H PBS buffer containing 17.5 nmole DSS-d6.1D ^2^H spectra were recorded on the ^2^H lock coil without locking, using an acquisition time of 0.3 s and a relaxation delay of 0.1 s, and 4096 transients (frequency drift < 0.2 Hz in 1 h acquisition). Free induction decays were zero-filled to 32768 data points and apodised with a 10 Hz exponential function, Fourier transformed, phased, baseline corrected and referenced to the internal DSS (natural abundance deuterium D-methylene resonances).

All NMR data were processed and analyzed using MestReNova software (MNova v 14, Santiago de Compostela, Spain). The ^1^H intensities in 1D ^1^H and 1D ^1^H-(^13^C)-HSQC spectra were manually integrated using the global spectra deconvolution (GSD) algorithm implemented in MNova software, which returns the area of each peak of interest. Then, the areas of each assigned and well-resolved metabolite from both ^1^H and ^1^H(^13^C)-HSQC spectra were normalized internally to the DSS peak at 0 ppm (integrated from 1D ^1^H spectra) and converted to the number of moles after accounting for the number of protons per group. The metabolite amounts and their isotopomers were normalized to protein content of the extracts. Metabolites and their isotopomers were assigned by comparing the obtained spectra with in-house multinuclear databases, public databases, or literature reports (Fan & Lane, 2008; Fan, 1996).

### Metabolism mass spectrometry

For the extracted metabolites, samples were homogenized in 80% methanol (v/v) containing internal standards (^13^C-labeled standards for quantification accuracy) at a ratio of 1 mL of solvent per 100 mg of tissue or 1x10^6^ cells. Homogenates were centrifuged at 14,000 g for 15 minutes at 4°C, and the supernatants were collected. The extracts were dried using a vacuum concentrator without heat and reconstituted in 100 µL of 50% methanol for mass spectrometry analysis.

For comprehensive metabolite analysis, both polar and non-polar metabolites were separated using different columns tailored for each metabolite class. Polar metabolites were analyzed using a Hydrophilic Interaction Liquid Chromatography (HILIC) column, which was maintained at 4°C, while non-polar metabolites were separated using a C18 reverse-phase column. Each column was optimized for their respective metabolite categories.

For the polar metabolites, mobile phase A consisted of water with 10 mM ammonium acetate and 0.1% formic acid, while mobile phase B was acetonitrile with 0.1% formic acid. For non-polar metabolites, an organic solvent mixture was used for the mobile phase, tailored for effective lipid extraction and separation. Chromatographic separation was achieved through gradient elution optimized for the respective metabolite classes. Data acquisition was carried out using multiple reaction monitoring (MRM) mode on a mass spectrometer with specific ion transitions tailored to each class of metabolites.

Mass spectrometric detection of metabolites from polar part was performed following the previously established method (Sun *et al*, 2021) by using a triple quadrupole mass spectrometer (e.g., AB Sciex QTRAP 6500 or similar), equipped with an electrospray ionization (ESI) source operating in both positive and negative ion modes. The ion source temperature was set at 450°C, with an ion spray voltage of 4500 V in positive mode and -4500 V in negative mode. The curtain gas was set at 35 psi, while both the nebulizer gas (GS1) and auxiliary gas (GS2) were set at 55 psi. Data acquisition was carried out in multiple reaction monitoring (MRM) mode, targeting specific m/z transitions for each metabolite of interest, optimized for the highest sensitivity and specificity. Polarity switching allowed the detection of both positive and negative ions in a single run, with stable isotope-labeled internal standards added to ensure accurate quantification across samples.

Mass spectrometry of lipids was performed using Direct Infusion Fourier Transform Mass spectrometry (DI-FTMS) as previously described (Fan *et al*, 2018). The ultra-high resolution (UHR)-FTMS raw data were assigned by our (CESB) in-house software PREMISE (PRecalculated Exact Mass Isotopologue Search Engine) that compares UHR-FTMS m/z data against our metabolite m/z library (calculated with mass accuracy to the 5th decimal point) to discern all known lipid MF and their 13C isotopologues, including hypothetical lipids, while simultaneously counting for the major adducts (here H+, Na+, K+ and NH4+). Natural abundance corrections were applied using an in-house algorithm. For statistical classification, we used only high accuracy monoisotopic m/z values that mapped to lipid molecular formulae, and multiple adducts of each were tracked throughout to avoid redundancy. 13C and 2H isotopologues of different lipid classes were evaluated from peak separations in the mass spectra, corresponding to different number of 13C, 2H and both present in the same compound (Lane *et al*, 2009).

Normalization was conducted based on protein concentration or cell number as appropriate. Metabolite concentrations were then compared across samples using statistical software, with significant differences determined by Student’s t-test or ANOVA, followed by post-hoc tests correcting for multiple comparisons.

Quality control samples, consisting of pooled sample extracts, were analyzed at regular intervals throughout the analytical run to monitor system performance, including retention time stability and response consistency. Additionally, blank samples (solvent only) were run to check for cross-contamination and background noise.

### DI-FTMS data analysis

DI-FTMS raw mass spectra were converted to mzML using the ThermoRawFileParser (Hulstaert *et al*, 2020). Scans were filtered to those of interest, and the raw intensity data was imported for each scan. Each scan was independently peak picked, and matching peaks determined across scans. Scans were normalized to a median scan. For each peak, the scans that contained non-zero intensities for that peak were determined, and the peak intensity and location was determined using all of the non-zero intensities across scans. Finally, scan level and aggregate level peak characteristics were exported in JSON for assignment by SMIRFE (Mitchell *et al*, 2019).

SMIRFE was used to generate assignments of the characterized peaks for each sample. Each sample was assigned with the appropriate labeling scheme, either C^13^, C^13^ + N^15^, or unlabeled. In all cases, the following adducts were allowed: H^+^, NH4^+^, K^+^, Na^+^. The caches were generated with the following limits: C: 130; N: 7; O: 28; P: 3; H: 230.

For each sample, the elemental molecular formula’s (EMFs) with an e-value score <= 0.1, and an M/Z <= 1400 was kept. The full set of EMFs were classified using the LipidClassifier (Mitchell *et al*, 2020). Those EMFs with a lipid or metabolite classification had their scores weighted two times higher than non-classified EMFs. M/Z deviation cutoffs were determined for each group of samples (C^13^-glucose labeled, C^13^-Ser/N^15^-Gln labeled, un-labeled) independently, and the set of M/Z cutoffs utilized for use during EMF voting of the entire set together.

For EMF voting, each sample has a group_EMF that is all the EMF assignments to a common set of peaks, as well as any EMFs that match the base formula but have different adducts. sudo_EMFs were created by iterative intersection - unions with other group_EMFs that share the same EMFs. The scores (either 1 - e-value or 2 * (1 - e-value)) of the EMFs across the samples were summed, and those EMFs within 90% of the max are kept. If any sample does not have matching EMFs, the peaks were checked to see if they match by both M/Z and relative intensity ordering.

Each isotopic molecular formula (IMF) within the EMF was then checked that the overall M/Z standard deviations were within the specified tolerances, if not individual peaks and potentially the whole IMF or EMF were removed, if the necessary peaks were not present. sudo_EMFs that share 50% or more of their peaks were merged, and EMF voting repeated.

To decide on majority lipid Categories, for each EMF any non-classified and non-lipid EMF assignments were removed, and then the remaining Categories were voted on. If there was a Category winner, that was reported as the EMF lipid Category. If the votes were tied, then the reported Category was “multiple”.

IMF level peaks were normalized using the median peak intensity for that sample. Missing values were imputed to a threshold value that is 2 times smaller than the smallest value generated by normalization in this case. IMF intensities were summed for each EMF, and then log2 transformed.

A linear model where the EMF intensity depends on whether the sample was PLK1 overexpressed or control was defined, with the labeling status (labeled, un-labeled) as well as group (1, 2, or 3) included as blocked variables. Analysis of variance (ANOVA) was run using the limma R package (Ritchie *et al*, 2015). EMFs with a raw p-value <= 0.1 were considered significantly changed.

EMF-EMF correlations were calculated using globally information theoretic weighted pairwise correlation, where Pearson correlations between EMFs are then weighted by information content and information consistency. Information content refers to how many entries are not missing in the two samples being correlated, and information consistency refers to how many entries are missing in common in the two samples being correlated.

EMF fold-changes were calculated as the average in PLK1 over the average in the controls. The minimum IMF M/Z across all samples is used for the EMF M/Z. EMFs with categories of “multiple”, “not lipid” and “not categorized” were removed.

Binomial statistics were calculated for each lipid category. EMFs with a log fold-change > 0 designated as positive, EMFs with a log fold-change <= 0 are designated as negative, and a binomial test is performed assuming a 50/50 split as the null statistic. P-values were adjusted using a Bonferroni correction.

### Graphical illustrations

Schematics of workflows were designed using Microsoft PowerPoint and Biorender.com.

### Statistical Analysis

Data were analyzed by GraphPad Prism 10 or Microsoft Excel unless otherwise stated. Data were modified by Photoshop 2023 or Microsoft PowerPoint unless otherwise stated. The significance of data sets was analyzed using the unpaired two-tailed t-test or Welch’s *t*-tests and p<0.05 was considered significant. Data are shown as means ± standard error of the mean (s.e.m) unless otherwise stated. n.s., not significant, * p<0.05, ** p<0.01, *** p<0.001, **** p<0.0001.

## Acknowledgements

We thank the Grant Support from National Institutes of Health grant R01 CA157429 (X. Liu), National Institutes of Health grant R01 CA196634 (X. Liu), National Institutes of Health grant R01 CA264652 (X. Liu), National Institutes of Health grant R01 CA256893 (X. Liu), University of Kentucky Markey Comprehensive Cancer Center for Cancer and Metabolism grant (COBRE) (X. Liu). Dr. Qing Shao provided the PHGDH protein simulation analysis. Dr. Ka Wing Fong helped us performed the gateway cloning for HA-PLK1. Dr. Eunus S. Ali provided constructive suggestions, comments and edits. Dr. Chi Wang helped with the bioinformatic analysis. These are all listed co-authors. Dr. Andrew Lane helped with NMR experiments and comments on the manuscript. Dr. Richard M. Higashi and Dr. Teresa W-M. Fan guided the MS experiments. We thank the University of Kentucky Center for Computational Sciences and Information Technology Services Research Computing for their support and use of the Lipscomb Compute Cluster and associated research computing resources. We also thank Markey Cancer Center Biospecimen Procurement and Translational Pathology Shared Resources Facilities (MCC BPTP SRF) for their support.

## Author contributions

X.R. initiated the project, performed the experiments, analyzed the data and wrote the manuscript. D.B.A. performed pathologic analysis of TMA slides and IHC slides. R.M.F. analyzed lipidomic data. P.L performed NMR. T.C and T.S performed MS. Y.W and D.H performed bioinformatic analysis. Y.Z and C.L provided tet-inducible plasmids and cell lines. X.W and J.P assisted immunoblotting. R.W and J.W performed the mouse study. Z.L and E.S.A provided suggestions and comments. Q.S performed simulation analysis. C. W, R.M.H, T. W-M. F, H.NB.M and X.L supervised the study. X.L led the whole process of the study. All authors commented the study and edited the manuscript.

## Expanded View Figure Legends

**Figure EV1. PLK1 is overexpressed in advanced prostate cancer**

(A). Expression of PHGDH across TCGA cancers (with tumor and normal samples).

(B). The expression of PLK1 in prostate cancer with Gleason Score 6, 7, 8, 9 and 10. Statistical analysis was performed using ANOVA.

(C). The survival curve of low and high PLK1 in TCGA-PRAD database. Statistical analysis was performed using Welch’s *t*-tests.

**Figure EV2. Expression level of PHGDH in cancer.**

Expression of PHGDH across TCGA cancers (with tumor and normal samples).

**Figure EV3. PHGDH is not regulated by AR**

(A). AR downregulation decreases the protein level of PHGDH. 22Rv1 cells (control and AR-KD) were subjected to IB.

(B). AR overexpression decreases the protein level of PHGDH. PC3 cells (control or AR-OE) were subjected to IB.

(C). PHGDH expression in the enzalutamide-resistant cells.

(D). PHGDH expression in the androgen deprived (ADT) and dihydrotestosterone (DHT)-treated LNCaP cells.

**Figure EV4. Preliminary simulation results of wild-type and PLK1-phosphorylated PHGDH in explicit solvents**

(A). The snapshots of the final confirmation of the wild-type (red) and phosphorylated (blue) PHGDH after a 1000 ns MD simulation.

(B). The radius of gyration of PHGDH-wt and PHGDH-S512-513-517Ph.

(C). Root mean square fluctuations (RMSF) of Cα atoms on the PHGDH.

(D). The secondary structure vs. simulation time for Residues 100-127.

(E). The secondary structure vs. simulation time for Residues 129-146.

(F). The secondary structure vs. simulation time for Residues 147-200.

**Figure EV5. PLK1 activation promotes ASCT2.**

(A). PLK1 overexpression (OE) promotes the protein level of ASCT2. A549 cells (WT or PLK1-OE) were subjected to immunoblotting (IB).

(B). Downregulation of PLK1 decreases the protein level of ASCT2 and constitutively active PLK1 (T210D) overexpression (TD) increases the protein level of ASCT2. U2OS cells (empty vector (EV), PLK1 knocking down (KD) or PLK1-TD) were subjected to immunoblotting (IB).

**Figure EV6. The mRNA level of serine transporters and enzymes involved in the sphingolipid metabolism.**

(A). The mRNA level of PHGDH, ASCT1, ASCT2, SPHK1, SPHK2, SPTLC1, SPTLC2, SPTLC3, SPTSSA after PLK1 overexpression in C4-2 cells.

(B). The mRNA level of PHGDH, ASCT1, ASCT2, SPHK1, SPHK2, SPTLC1, SPTLC2, SPTLC3, SPTSSA after PLK1 knockdown in DU145 cells.

**Figure EV7. The setup of stable cell lines.**

The setup of stable cell lines like C4-2-(shRNA Scramble, shRNA PHGDH 3’UTR, shRNA PHGDH 3’UTR-PHGDH WT, shRNA PHGDH 3’UTR-PHGDH 3D, shRNA PHGDH 3’UTR-PHGDH 3A)

**Figure EV8. The ^13^C incorporation into TCA cycle and PPP pathway.**

(A). The comparison of ^13^C incorporation into TCA cycle between C4-2-2D/C4-2-WT and LNCaP-PLK1/LNCaP-control cells.

(B). The comparison of ^13^C incorporation into PPP pathway between C4-2-2D/C4-2-WT and LNCaP-PLK1/LNCaP-control cells.

Statistical analysis was performed using unpaired two-tailed *t*-tests (A-B).

**Figure EV9. The ^13^C incorporation into nucleotide pathways and other glucose-related metabolic pathways.**

(A). The comparison of ^13^C incorporation into nucleotide synthesis between C4-2-2D/C4-2-WT and LNCaP-PLK1/LNCaP-control cells.

(B) and (C). The comparison of ^13^C incorporation into glucose-related metabolites between C4-2-2D/C4-2-WT and LNCaP-PLK1/LNCaP-control cells.

Statistical analysis was performed using unpaired two-tailed *t*-tests (A-C).

**Figure EV10. The ^2^H incorporation into the downstream metabolites of serine metabolism.**

(A). The flow chart of ^2^H incorporation into the downstream metabolites of serine metabolism.

(B). The ^2^H incorporation into the downstream metabolites of serine metabolism from LNCaP-PLK1 and LNCaP-control cells.

(C). The ^2^H incorporation into the downstream metabolites of serine metabolism from C4-2-2D and C4-2-WT cells.

(D). The comparison of thymidine labelling (dTDP, dTMP) with ^2^H from [2,3,3-2H]-serine between LNCaP-PLK1 vs LNCaP-Control cells and C4-2-PHGDH-3D vs C4-2-PHGDH-WT cells.

Statistical analysis was performed using unpaired two-tailed *t*-tests (B-D).

**Figure EV11. The NMR analysis for the media and polar parts of WT, 3D and 3A C4-2 cells.**

(A). The 1D ^1^H(^13^C)-HSQC NMR analysis for the glucose consumption, pyruvate production and lactate production for LNCaP-PLK1 and LNCaP-control cells. Statistical analysis was performed using the two-way ANOVA.

(B). The 1D ^1^H(^13^C)-HSQC NMR analytic comparations for the glucose consumption and lactate production for C4-2-3A and C4-2-WT cells.

(C). The 1D ^1^H(^13^C)-HSQC NMR analytic comparation for the glucose consumption and lactate production for C4-2-3A and C4-2-3D cells.

(D). The 1D ^1^H(^13^C)-HSQC NMR analysis for the glutamine, alanine and glutamate from media of C4-2-3A and C4-2-WT cells.

(E). The 1D ^1^H(^13^C)-HSQC NMR analysis for polar part of C4-2-3A and C4-2-WT cells. Statistical analysis was performed using Welch’s t-test (B-F).

**Figure EV12. The gene heatmap of serine related KEGG metabolic pathway in C4-2 WT and 3A cells.**

(A). The analysis of gene heatmap and the KEGG pathways of sphingolipid metabolism for C4-2 WT and 3A cells.

(B). The analysis of gene heatmap and the KEGG pathways of glycine, serine and threonine metabolism for C4-2 WT and 3A cells.

**Table EV1. The samples information of the lipidomic analysis for the LNCaP-control and LNCaP-PLK1 cells.**

**Table EV2: Mass Spectrum analysis for the phosphorylation of PHGDH peptides.**

**Table EV3. The proportions of ^2^H labeled lipid species among the grand total lipids.**

**Table EV4. The proportions of ^2^H labeled sphingolipid species.**

